# DPYSL3B is a regulator of chemoresistance via DNA repair and metabolic reprogramming in prostate cancer

**DOI:** 10.1101/2025.07.03.658029

**Authors:** Roosa Kaarijärvi, Heidi Kaljunen, Nella Itkonen, Kseniia Bureiko, Kirsi Ketola

## Abstract

Prostate cancer is among the most common cancers worldwide. Aggressive, treatment-resistant subtypes including androgen receptor (AR) -negative and neuroendocrine prostate cancer, present a significant clinical challenge. Platinum-based chemotherapies, such as carboplatin, remain a treatment option for these subtypes, but their efficacy is limited due to activation of DNA damage response (DDR) pathways.

Here, we report that DPYSL3 (CRMP4) is highly expressed in AR-negative prostate cancer, especially in liver metastases. We show that prostate cancer cells express two transcript variants of DPYSL3, DPYSL3A and DPYSL3B. While DPYSL3A has tumor-suppressive roles, the function of DPYSL3B remains unclear.

In this study, we demonstrate that high DPYSL3B expression correlates with poor patient survival in prostate cancer. Functional studies revealed that DPYSL3B silencing impairs proliferation and sensitizes prostate cancer cells to carboplatin-induced DNA damage. Gene set enrichment analyses of DPYSL3B-depleted cells showed negative enrichment of pathways related to the cell cycle and DNA repair, including E2F1, AURKA, homologous recombination, and the Fanconi anemia (FA) pathway, consistent with impaired DNA repair capacity. This was accompanied by γH2AX foci accumulation and apoptosis, as indicated by increased caspase-3/7 activity, particularly under carboplatin exposure. Metabolomic profiling further revealed decreased levels of glutathione and nucleotide precursors, suggesting a role for DPYSL3B in redox homeostasis and nucleotide biosynthesis.

Taken together, our transcriptomic, metabolomic, and functional analyses identify DPYSL3B as a regulator of DNA repair and metabolic reprogramming and support its potential as a therapeutic target to overcome chemoresistance in aggressive, treatment-resistant AR-negative prostate cancer.

## Introduction

Prostate cancer (PCa) is the second most commonly diagnosed cancer in men worldwide, and the number of new cases annually is projected to double by 2040^1,2^. PCa is also the second leading cause of cancer-related mortality among men. The development of aggressive and treatment-resistant subtypes, such as androgen receptor (AR) -negative and neuroendocrine prostate cancer (NEPC), is a major contributor to poor prognosis^1,3^. Currently, treatment options for advanced and aggressive metastatic PCa are largely limited to platinum-based chemotherapies, which provide only short-term clinical benefit^4^. These agents, including carboplatin, cisplatin, and oxaliplatin, bind to DNA and cause DNA damage by inducing interstrand crosslinks (ICLs), thereby impairing replication^5^. This, in turn, triggers the DNA damage response (DDR) and the recruitment of DNA repair proteins^5^. Cancer cells exploit DDR-associated pathways and proteins to evade treatment, and thus detailed understanding of these pathways and their role in treatment resistance is essential^6^.

We have recently identified DPYSL5 (Dihydropyrimidinase Like 5, also known as Collapsin Response mediator protein 5 or CRMP5) as highly expressed in treatment-induced NEPC (t-NEPC), where it promotes lineage plasticity via PRC2/EZH2^7^. In this study, we investigated whether other DPYSL family members are implicated in aggressive prostate cancer. Patient transcriptomic data revealed that DPYSL3 (also known as CRMP4) is highly expressed in double-negative prostate cancer (DNPC) compared to other PCa subtypes. DPYSL3 expression has been linked to poor prognosis in several cancers, including lung cancer, urothelial carcinoma, thyroid carcinoma, neuroblastoma, renal cell carcinoma, gastric cancer, and liver cancer^8–14^. Conversely, some studies suggest tumor-suppressive functions for DPYSL3 in lung and prostate cancer^15,16^. These opposing roles may be explained by the existence of two distinct splice variants, DPYSL3A and DPYSL3B (known also as CRMP4a and CRMP4b), which have been proposed to exhibit opposite functions in cancer biology^17^. A similar phenomenon has been described in neuronal development, where DPYSL3B localizes to the growth cone and promotes neurite extension, whereas DPYSL3A is diffusely distributed and has no effect on neuronal outgrowth when overexpressed^18^. In the context of cancer, this functional variance remains largely unexplored, as most studies fail to distinguish between DPYSL3 isoforms.

In prostate cancer, prior research on DPYSL3 has primarily focused on the DPYSL3A/CRMP4a isoform, considered a putative tumor suppressor^15,19,20^. Overexpression of DPYSL3A has been shown to inhibit cell migration and invasion^21^. Mechanistically, DPYSL3A is suggested to participate in cytoskeletal organization through interacting with RhoA and suppressing its function resulting in reduced cell spreading^19,21^. In contrast, the role of the DPYSL3B isoform in prostate cancer remains largely uncharacterized.

Here, we show that high expression of the DPYSL3B isoform is associated with reduced overall survival in PCa patients. Moreover, DPYSL3 expression is upregulated in response to enzalutamide, a second-generation antiandrogen used in androgen-deprivation therapy (ADT) for castration-resistant prostate cancer. Functionally, DPYSL3B silencing reduces PCa cell proliferation, and downregulates gene sets associated with cell cycle regulation and cancer progression. In addition, our findings reveal that DPYSL3B-depleted prostate cancer cells show increased sensitivity to carboplatin, resulting in increased accumulation of DNA damage. In our previous work, we showed that DNA damage-responsive FANCI and the associated Fanconi anemia (FA) pathway contribute to carboplatin resistance in prostate cancer^22,23^. Here, we show that DPYSL3B depletion leads to downregulation of FA pathway, suggesting a role for DPYSL3B in regulating FA pathway-mediated DNA damage repair in prostate cancer. Furthermore, metabolomic profiling of DPYSL3B-silenced cells revealed reduced levels of glutathione and nucleotide precursors, suggesting that DPYSL3B may also contribute to redox and metabolic homeostasis in prostate cancer cells.

## Results

### DPYSL3 is highly expressed in AR-negative prostate cancer and enriched in liver metastases

We have previously shown that the expression of DPYSL5, a member of DPYSL protein family, is induced under long-term androgen deprivation and promotes lineage plasticity and treatment resistance in prostate cancer^7^. Here, we investigated whether other members of the DPYSL family, specifically DPYSL2, DPYSL3 and DPYSL4, might similarly contribute to prostate cancer aggressiveness and treatment resistance. To assess whether prolonged AR inhibition affects the expression of these DPYSL family members, we exposed LNCaP prostate cancer cells to the AR inhibitor enzalutamide (ENZ) for 9 days and analysed the DPYSL2-4 mRNA levels using RT-qPCR. The expression of all three DPYSL transcripts was significantly increased by day 9 of ENZ (Fig. 1a). These results indicate that AR inhibition induces the expression of DPYSL family members and raises the possibility that also other DPYSL family members, in addition to DPYSL5, may contribute to treatment resistance mechanisms in prostate cancer.

**Figure 1.**
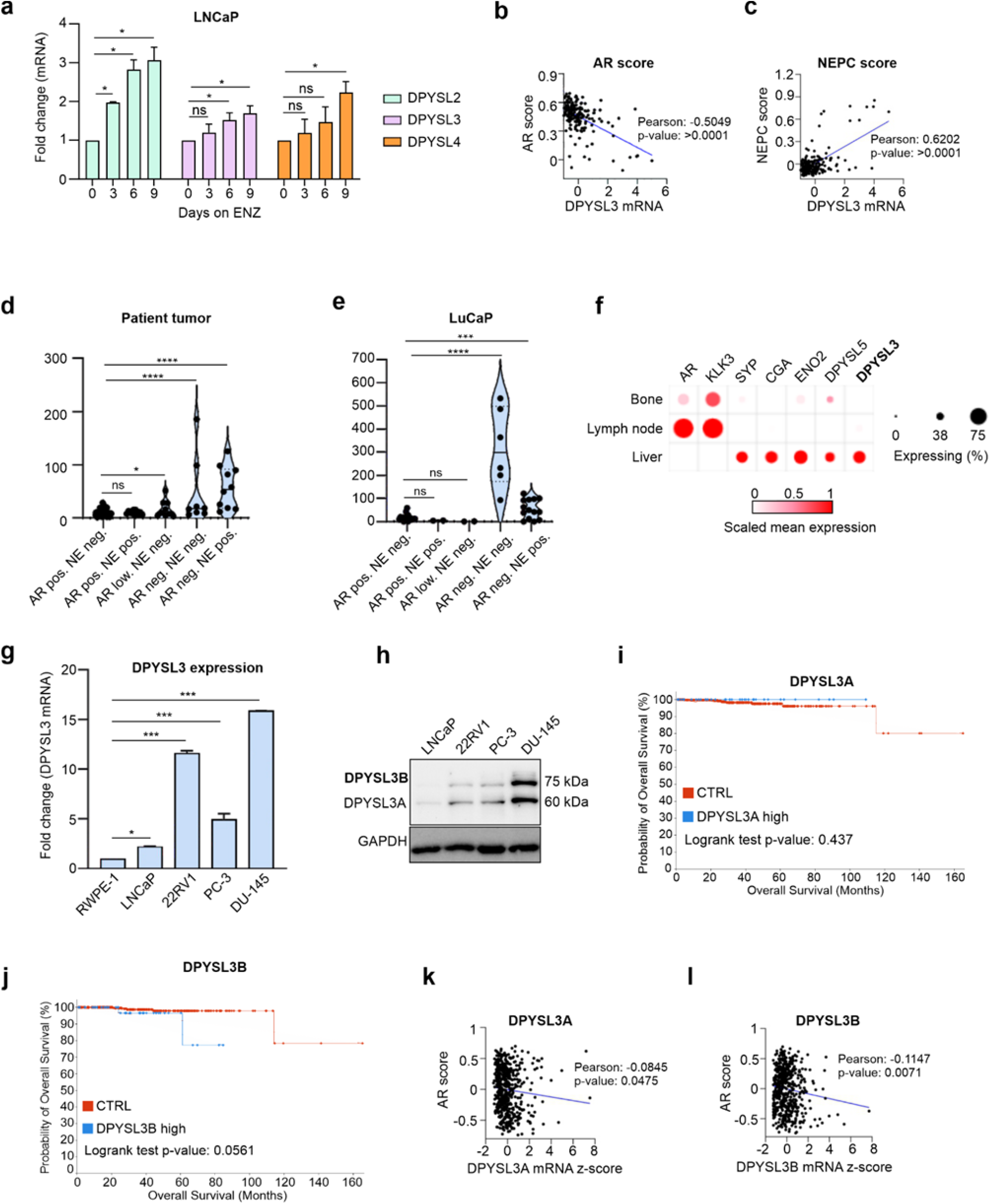
High DPYSL3 expression correlates with AR-negative prostate cancer subtype. a) DPYSL2-4 mRNA expression in response to enzalutamide (ENZ) exposure in LNCaP cells analysed using RT-qPCR. All genes were significantly upregulated after 9 days of ENZ exposure. Bars represent mean ± SD with n = 3. P-values shown as asterisks (*p≤ 0.05, **p ≤ 0.01 and ***p ≤ 0.001). b) DPYSL3 expression correlates negatively with AR score, and c) positively with NEPC score based on analysis of the Abida *et al*. 2019 dataset. d) High DPYSL3 expression was observed in the double-negative (AR- and NE- negative) PCa subtype, based on analysis of the Labrecque *et al*. 2019 dataset e) High DPYSL3 expression in double-negative PCa subtypes in LuCaP PCa patient-derived xenografts. f) DPYSL3 and NE-marker expression (DPYSL5, ENO2/NSE, SYP and CGA)and AR-markers (AR, KLK3/PSA) in single-cell transcriptomic data from three distinct metastatic PCa sites. High DPYSL3 expression was seen in the liver metastases, which also had high NE marker expression. Conversely, in AR positive lymph node metastases expression of DPYSL3 was low. DPYSL3 expression thus correlates with increased disease aggressiveness. g) DPYSL3 expression was determined on mRNA level in RWPE-1 and in both AR positive (LNCaP and 22Rv1) and AR negative (PC-3 and DU-145) prostate cancer cell lines using RT-qPCR. The highest DPYSL3 expression was seen in DU-145 cell line. Bars represent mean±SD with n=3. P-values shown as asterisks (*p≤ 0.05, **p ≤ 0.01 and ***p ≤ 0.001). h) Protein levels of DPYSL3 analysed in prostate cancer cell lines. Highest expression was observed in AR negative cell lines. Additionally, expression of two different isoforms, DPYSL3A (∼60 kDa) and B (∼75 kDa), were observed in all cell lines. i) The effect of DPYSL3 isoforms on overall patient survival was analysed using Taylor et al. prostate cancer patient dataset. High expression of DPYSL3A isoform had no effect on survival while j) high DPYSL3B expression correlated with poor overall survival. Even though both k) DPYSL3A and l) DPYSL3B expression correlated negatively with AR score, the correlation was significantly higher with DPYSL3B.

Due to the observed increase in DPYSL mRNA levels in response to ENZ in PCa cell lines, we next investigated whether DPYSL mRNA expressions correlate with prostate cancer aggressiveness in clinical datasets. To this end, we analysed the dataset from Abida *et al*. 2019 and found that DPYSL3 expression negatively correlated with AR score (Fig. 1b) and positively correlated with NEPC score (Fig. 1c). A similar strong correlation was not observed with DPYSL2 or DPYSL4 (Supplementary Fig. S1 a and b). While DPYSL3 mRNA was detected across all PCa subtypes, it was significantly upregulated in the double-negative subtype (lacking both AR and neuroendocrine markers) in the Labrecque *et al*. 2019 dataset (Fig. 1d). A similar pattern was observed in LuCaP prostate cancer patient-derived xenografts, where DPYSL3 expression was highest in double-negative tumors (Fig. 1e). In addition, we analysed the expression of DPYSL3 alongside known NE markers (DPYSL5, ENO2/NSE, SYP and CGA) and AR-markers (AR, KLK3/PSA) in single-cell transcriptomic data from three distinct metastatic PCa sites (Fig. 1f)^24,25^. Lymph node metastases showed high AR and KLK3 expression, whereas liver metastases expressed DPYSL3 and NE markers. Notably, liver metastases in PCa are often associated with more aggressive, AR-independent disease, while lymph node metastases typically retain AR signaling and have better prognosis^26^. These findings suggest that DPYSL3 expression is elevated in AR-negative, clinically aggressive PCa subtypes.

To further confirm these associations in PCa cells, we quantified DPYSL3 mRNA levels in AR-positive (LNCaP and 22Rv1) and AR-negative (PC-3 and DU-145) using RT-qPCR. Benign RWPE-1 prostate epithelial cells were used as a reference. DPYSL3 mRNA levels were highest in AR-negative DU-145 cells (Fig. 1g). Protein-level analysis by western blot confirmed these results, with strongest DPYSL3 protein expression in AR-negative PC-3 and DU-145 cells (Fig. 1h). Interestingly, two distinct bands were consistently detected across all PCa cells, corresponding to the predicted molecular weights of the two known DPYSL3 isoforms: DPYSL3A (∼60 kDa, Fig. 1g) and DPYSL3B (∼75 kDa, Fig. 1h). This observation prompted us to further investigate the expression patterns and potential functions of these isoforms individually, utilizing publicly available PCa patient datasets.

### High DPYSL3B expression correlates with poor overall survival in prostate cancer patients

It has previously been suggested that the DPYSL3A and B isoforms have opposing functions, with DPYSL3A acting as a tumor suppressor and DPYSL3B functioning as an oncogene in gastric cancer^27^. We therefore investigated whether these variants exert differential effects on overall survival in prostate cancer patients. To this end, we first analysed the expression of DPYSL3 variants in the Taylor *et al*. PCa dataset. Interestingly, DPYSL3A expression did not affect overall survival (Fig. 1i) whereas patients with high DPYSL3B expression had significantly poorer overall survival (Fig. 1j). Moreover, while the expression of both A and B variants negatively correlated with AR score (Fig. 1k and 1l), the correlation was stronger for DPYSL3B (Fig. 1l). Taken together, these results suggest that the DPYSL3B isoform may contribute to PCa progression and was therefore selected for further investigation.

### DPYSL3B silencing reduces the proliferation of prostate cancer cells

Next, we investigated the potential pro-oncogenic role of DPYSL3B in prostate cancer cells using DPYSL3B silencing. First, we assessed whether DPYSL3B silencing affects cell proliferation in high DPYSL3 protein expressing PC-3 and DU-145 cells (Fig. 1g). A small interfering RNA (siRNA) targeting specifically DPYSL3B variant was used, and cell proliferation was monitored using the IncuCyte S3 live-cell imaging system and its associated software. LNCaP cells, which exhibited the lowest DPYSL3 expression among the cell lines studied (Fig. 1g) were included for comparison. The results showed a significant reduction in cell proliferation (p-value <0,01) in response to DPYSL3B siRNA in all three cell lines compared to non-targeting siRNA control (Fig. 2a-c), indicating that DPYSL3B is important for PCa cell survival.

**Figure 2.**
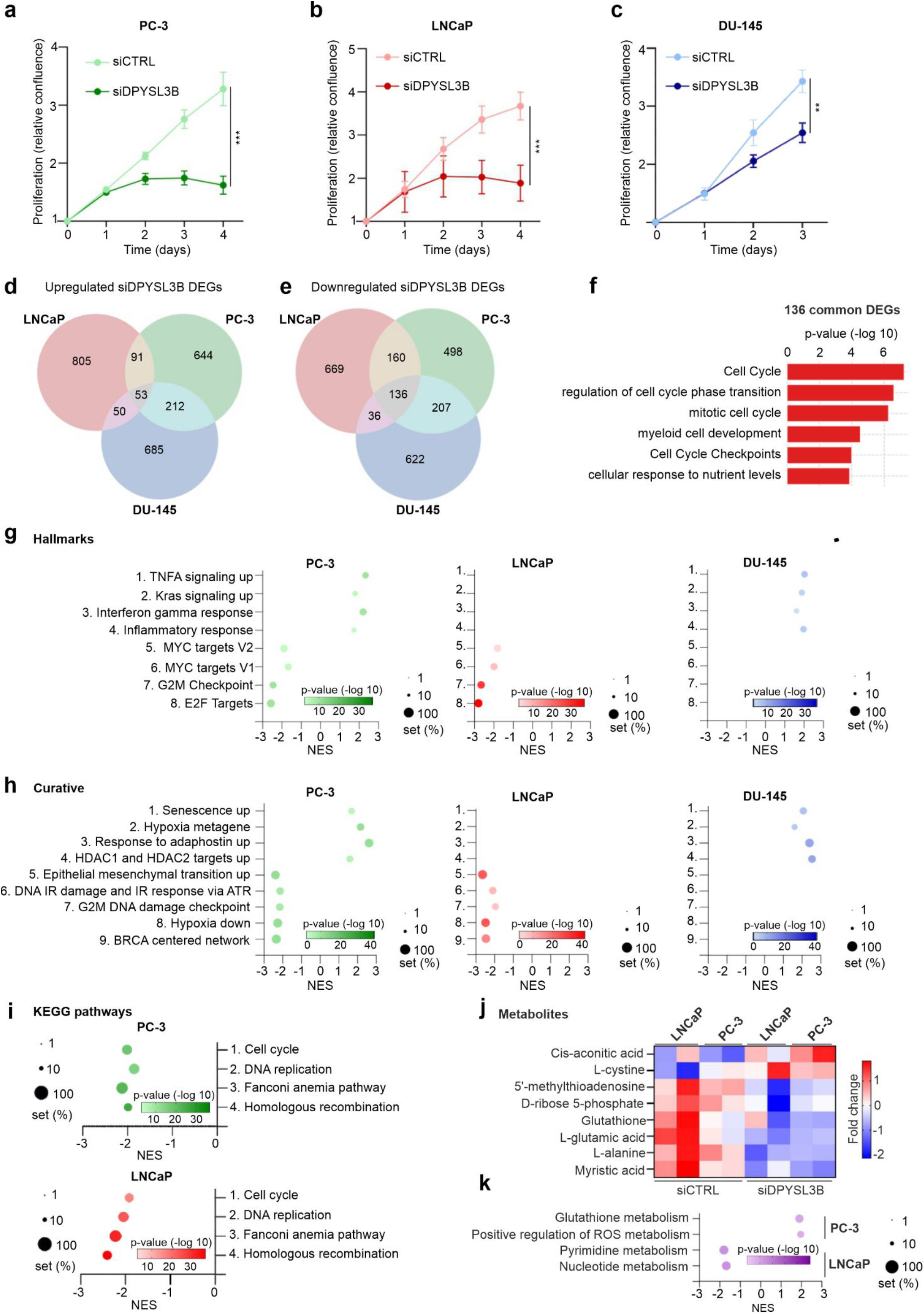
DPYSL3B depletion reduces proliferation and downregulates DNA repair-related pathways. Proliferation of a) PC-3 cells, b) LNCaP cells and c) DU-145 cells in response to DPYSL3B silencing with siRNA (siDPYSL3B) was studied using IncuCyte S3 live-cell imaging system. Non-targeting siRNA (siCTRL) was used as control. DPYSL3B depletion significantly reduced proliferation when compared to control siRNA in all cell lines. Bars represent mean ± SD with n = 4. p-values shown as asterisks (*p ≤ 0.05, **p ≤ 0.01 and ***p ≤ 0.001) d) Comparison of top 1000 upregulated DEGs in LNCaP, PC-3 and DU-145 cells in response to DPYSL3B silencing with siRNA. 53 DEGs were common between all cell lines. e) The top 1000 downregulated DEGs in LNCaP, PC-3 and DU-145 cells in response to DPYSL3B siRNA were compared. 136 DEGs were common between all cell lines. f) Gene set enrichment analysis (GSEA) of the 136 common downregulated DEGs indicated enrichment in pathways related to cell cycle, myeloid cell development and nutrient sensing. g) GSEA (Hallmark gen sets) in PC-3, LNCaP and DU-145 cells in response to DPYSL3B depletion. PC-3 cells showed positive enrichment of TNFA signaling, Kras signaling, interferon gamma response and inflammatory response; and negative enrichment of Myc targets V2 and V1, G2/M checkpoint, and E2F targets. LNCaP cells shared negatively enriched gene sets with PC-3 cells while lacked positive enrichment. Conversely, DU-145 cells shared positively enriched gene sets with PC-3 cells whereas no negatively enriched gene sets were found. (n = 3, adjusted p-value < 0.05 and false discovery rate (FDR) <0.25) h) GSEA of curative gene sets in response to DPYSL3B depletion. In PC-3 cells, positively enriched genes were related to senescence, genes upregulated in response to hypoxia and adaphostin, HDAC1 and HDAC2 targets. Negatively enriched genes in PC-3 cells were related to epithelial-mesenchymal transition, DNA IR damage, G2/M DNA damage checkpoint, genes downregulated in response to hypoxia and BRCA centered network. LNCaP cells shared negatively enriched gene sets with PC-3 cells while no positively enriched sets were found. Conversely, DU-145 cells shared positively enriched gene sets with PC-3 cells. i) KEGG pathway analysis showed that DPYSL3B silenced PC-3 and LNCaP cells led to negative enrichment of pathways involved in cell cycle, DNA replication, Fanconi anemia, and homologous recombination. j) Metabolomics profiling of DPYSL3B-silenced PC-3 and LNCaP cells revealed reduced levels of metabolites essential for nucleotide and protein synthesis (e.g. D-ribose 5-phosphate, L-glutamic acid), and increased levels of L-cystine, consistent with increased oxidative stress. Cells exposed to non-targeting siRNA (siCTRL) were used as controls. k) GSEA focusing on metabolism-related gene sets revealed positive enrichment of genes related to glutathione and ROS metabolism in PC-3 cells, and negative enrichment of genes related to pyrimidine and nucleotide metabolism in LNCaP cells.

To further understand the effect of DPYSL3B silencing on molecular level in the selected cell lines (LNCaP, PC-3 and DU-145), we performed RNA-sequencing using non-targeting siRNA exposed cells as control. Using the obtained RNA-seq data, we first investigated the top 1000 up-and downregulated differentially expressed genes (DEGs) that were common among the three cell lines (Supplementary Table S1). Surprisingly, though there were 53 commonly upregulated genes, they showed no enrichment in any pre-defined gene sets (Fig. 2d). However, when comparing the downregulated DEGs common between the three cell lines, we discovered enrichment of genes related to regulation of cell cycle and cell cycle checkpoints, such as AURKA and MYBL2, which both have been reported to be highly expressed in aggressive PCa variants^28,29^, and also enrichment of associated gene sets (Fig. 2e and 2f). These findings suggest a potential role for DPYSL3B in regulating the cell cycle in PCa cells.

### DPYSL3B silencing alters the expression of genes involved in inflammation, hypoxia, and DNA damage

The investigation of RNA-seq data from DPYSL3B silenced LNCaP, PC-3 and DU-145 cells revealed that there were commonly up-and downregulated genes among the three cell lines. We next investigated the pathways affected by DPYSL3B silencing in each cell line separately in more detail. To this end, we performed a gene set enrichment analysis (GSEA) on the RNA-seq data from DPYSL3B silenced cells. Interestingly, when looking into hallmark gene sets, we observed a different response to DPYSL3B silencing between the cell lines. More specifically, while in PC-3 cells we detected both positively and negatively enriched gene sets (Fig. 2g, left panel), in LNCaP, only negatively enriched gene sets (Fig. 2g, middle panel) and in DU-145 cells, only positively enriched gene sets were observed (Fig. 2g, right panel). A more detailed investigation of these gene sets revealed that genes related to inflammation (TNFA signaling, Kras signaling and interferon gamma response) were positively enriched in both PC-3 and DU-145, indicating cellular stress response activation upon DPYSL3B depletion in these cell lines (Fig. 2g). In contrast, gene sets related to cell cycle and cancer progression, such as G2/M checkpoint, E2F and MYC targets, were negatively enriched in PC-3 but not in DU-145 cells, suggesting that PC-3 cells are more dependent on DPYSL3B in their survival when compared to DU-145 (Fig. 2g). In LNCaP cells, DPYSL3 depletion did not induce inflammation related pathways but like in PC-3 cells, cell cycle and cancer progression related gene sets were negatively enriched (Fig. 2g, middle panel). Additionally, previous research of DPYSL3A on PC-3 cells has reported DPYSL3A regulating expression of E2F1^20^. We utilized Enrichr and our RNA-seq data to discover which transcription factors are known to regulate the downregulated DEGs in response to DPYSL3B depletion^30–32^. We indeed detected a strong enrichment in E2F1 targets in all our cell lines (Supplementary Fig. S2a) and downregulation of E2F1 itself (Supplementary Fig. 2b).

Next, we investigated the curated gene sets from GSEA to further analyze the response to DPYSL3B depletion in the three cell lines. Similar pattern as with curated gene sets were observed, where in PC-3 cells there were both positively and negatively enriched gene sets (Fig. 2h, left panel) while in LNCaP cells there was only negatively (Fig. 2h, middle panel), and in DU-145 cells only positively enriched gene sets (Fig. 2h, right panel). In both LNCaP and PC-3 cells an enrichment of DNA damage and hypoxia related gene sets was observed among the downregulated genes (Fig. 2h, left panel). More specifically, these included genes typically suppressed under hypoxia (MANALO_HYPOXIA_DN), genes related to the G2/M DNA damage checkpoint (REACTOME_G2_M_DNA_DAMAGE_CHECKPOINT), genes involved in BRCA centered network (PUJANA_BRCA_CENTERED_NETWORK) and genes involved in responding to DNA damage caused by IR (ionizing radiation) (WP_DNA_IRDAMAGE_AND_CELLULAR_RESPONSE_VIA_ATR). Additionally, when we analysed enriched KEGG-pathways, we found that in both LNCaP and PC-3 cells genes related to cell cycle, DNA replication and interestingly Fanconi anemia (FA) pathway were downregulated (Fig. 2i). Furthermore, we detected downregulation of homologous recombination related genes (Fig. 2i) which are required for repairing DNA double strand breaks and include members of FA pathway^33^. FA-pathway is a well-known part of DNA damage response and is particularly important in cancer therapy context via its ability to repair replication hindering interstrand crosslinks induced by platinum-based chemotherapy reagents, such as carboplatin^23^.

Interestingly, when we inspected the positively enriched curated gene sets in response to DPYSL3B depletion in both PC-3 and DU-145 cells (Fig. 2h, left and right panel, respectively), we found gene sets associated with hypoxia or oxidative stress. These included gene set upregulated in response to hypoxia (WINTER_HYPOXIA_METAGENE), genes upregulated after HDAC1 and HDAC2 silencing (SENESE_HDAC1_AND_HDAC2_TARGETS_UP), genes upregulated after adaphostin (tyrosine kinase inhibitor^34^) treatment (PODAR_RESPONSE_TO_ADAPHOSTIN_UP) and senescence related gene set (FRIDMAN_SENESCENCE_UP). HDAC1/2 silencing and adaphostin treatment have been shown to promote apoptosis and oxidative stress^35,36^ while hypoxia has been shown to promote senescence^37^. Overall, DPYSL3B depletion in both PC-3 and DU-145 cells leads to oxidative stress responses indicating that DPYSL3B is involved in hypoxia regulation.

Taken together, our RNA-sequencing data showed differential responses to DPYSL3B depletion in LNCaP, PC-3 and DU-145 cell lines. In DU-145 cells upregulation of both inflammation and oxidative stress related genes and enrichment of associated gene sets were observed upon DPYSL3B silencing, indicating increased cellular stress response. A similar response to DPYSL3B depletion was seen in PC-3 cells, with the additional negative enrichment of cancer progression, cell cycle regulation and DNA damage related gene sets. In LNCaP cells the same gene sets were downregulated as in PC-3 cells. Overall, these results suggest that DPYSL3B supports PCa cell survival, potentially by regulating cell cycle progression and the DNA damage response.

### DPYSL3B depletion disrupts antioxidant metabolism

As our RNA-seq analysis and GSEA indicated that DPYSL3B is involved in key cellular stress responses, we next examined how DPYSL3B depletion affects cellular metabolism. For this, LNCaP and PC-3 cells were transfected with DPYSL3B-targeting siRNA (siDPYSL3B) or non-targeting control siRNA (siCTRL), and subjected to metabolic profiling by LC-MS. We detected over 150 metabolites (Supplementary Table 2) and compared the common down-and upregulated metabolites between siDPYSL3B and siCTRL conditions in both cell lines (Fig. 2j). Interestingly, we observed upregulation of L-cystine and downregulation of L-glutamic acid (glutamate) and glutathione (GSH). L-cystine, the oxidized dimer form of cysteine, is normally reduced to cysteine upon cellular uptake^38^, whereas GSH is a key antioxidant that protects cells from oxidative stress^38^. Both glutamate and cysteine are essential substrates GSH synthesis; therefore lack of glutamate may impair GSH production^38^. The concurrent accumulation of L-cystine and reduction of glutamate and GSH levels in both LNCaP and PC-3 cells in response to DPYSL3B silencing thus indicates increased oxidative stress, in line with our GSEA findings.

Additionally, we observed downregulation of several metabolites important for cellular proliferation and homeostasis, including D-ribose 5-phosphate, which is required for nucleotide synthesis, amino acid L-alanine, 5’-methylthioadenosine, which is a nucleoside involved in modulating inflammation, and myristic acid, which is required for protein N-myristoylation, a process that affects subcellular tracking and localization, and thus function of several oncogenic proteins^39–41^. To further support these finsings, we re-examined our GSEA results focusing on metabolism-related gene sets. In PC-3 cells, glutathione and ROS metabolism pathways were upregulated, while in LNCaP cells, pyrimidine and nucleotide metabolism pathways were downregulated (Fig. 2k). Taken together, these results indicate that DPYSL3B depletion disrupts antioxidant metabolism and impairs nucleotide biosynthesis, two processed critical for the survival of rapidly proliferating cancer cells.

### DPYSL3B depletion leads to downregulation of the Fanconi anemia DNA repair pathway

Our RNA-sequencing and metabolomics analyses indicated that DPYSL3 affects oxidative stress responses and DNA damage repair, particularly Fanconi anemia pathway. To explore the potential role of DPYSL3B in regulating of FA pathway, we first composed a FANCI up- and downregulated signature gene signature from the common DEGs between FANCI depleted LNCaP and PC-3 cells using our published RNA-seq data^23^. We then compared this FANCI signature set to all DEGs in DPYSL3B silenced PC-3, LNCaP and DU-145 cells and found an enrichment of FANCI targets (Fig. 3a). We then further analysed the expression of other FA pathway genes using RNA-seq data from DPYSL3B silenced PC-3, LNCaP and DU-145 cells and detected downregulation of several FA complex members (Fig. 3c). This suggests that DPYSL3 can regulate FA pathway and thus the related DNA damage repair.

**Figure 3.**
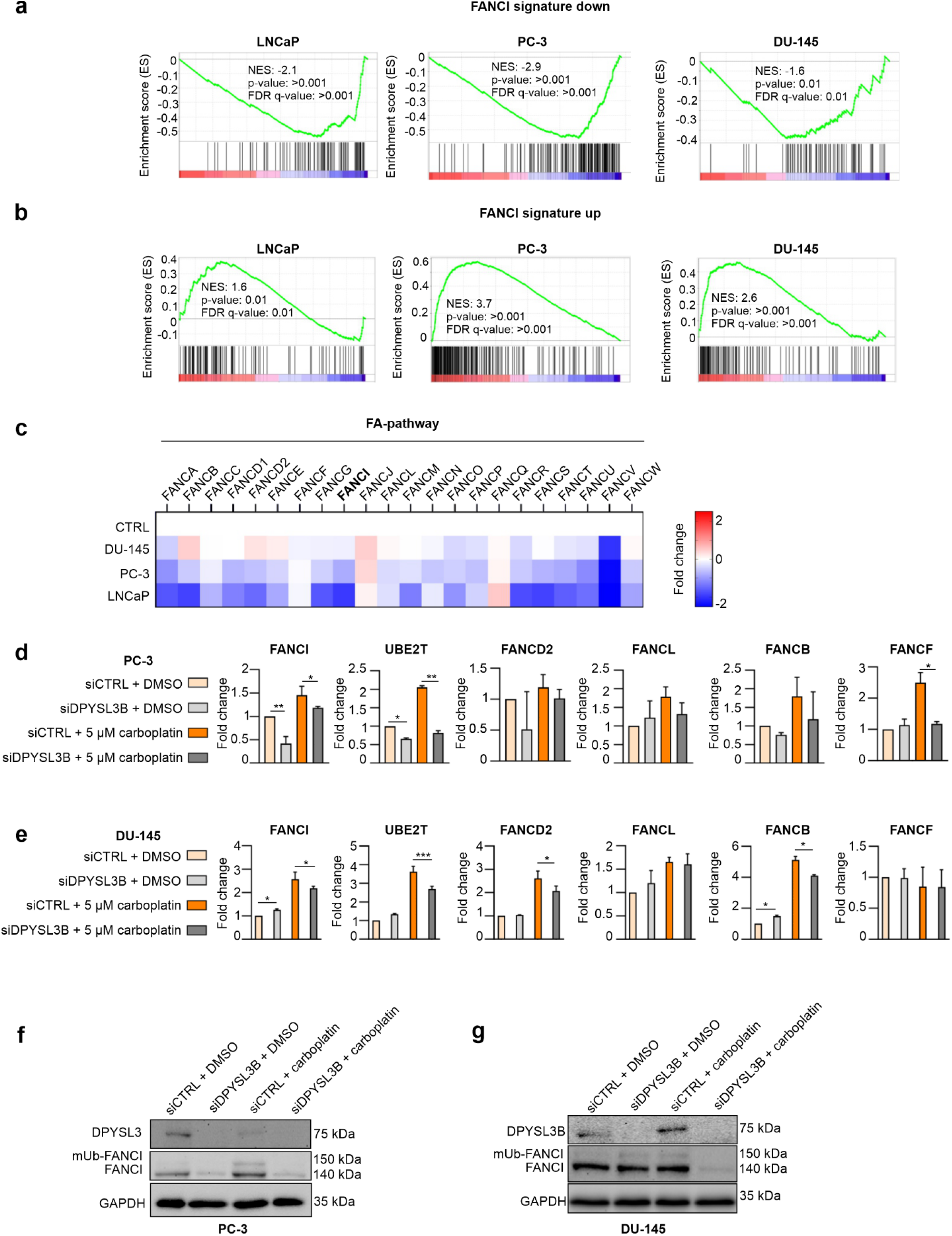
DPYSL3B depletion downregulates Fanconi anemia complex members. a) Comparison of DPYSL3 siRNA data from LNCaP, PC-3 and DU-145 cells to our FANCI signature set consisting of genes enriched by FANCI depletion (siFANCI) (Kaljunen et al. 2023) showed enrichment of FANCI targets upon DPYSL3B depletion. Genes that were downregulated by siFANCI were also downregulated by DPYSL3B silencing and b) conversely, genes that were upregulated by siFANCI were upregulated by DPYSL3B. c) The analysis of DPYSL3B silencing RNA-seq data showed that DPYSL3B depletion led to downregulation of FA complex members, including FANCI, which is the main regulator of this complex. The results are presented as a heatmap with fold change compared to the corresponding control for each cell line d) The mRNA expression of FA complex members was analysed using RT-qPCR in PC-3 and in e) DU-145 cells exposed to DPYSL3B silencing with or without DNA damage inducing and FA complex activating carboplatin. Non-targeting siRNA and DMSO were used as controls. The results showed that DPYSL3B silencing leads to downregulation of FA complex members also in the presence of carboplatin. Bars represent mean±SD with n=3. P-values shown as asterisks (*p≤ 0.05, **p ≤ 0.01 and ***p ≤ 0.001). f) Western blot analysis was conducted from DPYSL3B (siDPYSL3) or non-targeting siRNA (siCTRL) exposed PC-3 or g) DU-145 cells with DMSO or carboplatin to analyse FANCI protein levels. Monoubiquitinated FANCI (mUb-FANCI), denoting the active form of FANCI, was observed in siCTRL samples upon exposure to carboplatin. In both PC-3 and DU-145 cell lines combined exposure to DPYSL3B siRNA and carboplatin resulted in depletion of mUb-FANCI and FANCI itself.

FA pathway is known to repair ICL type DNA damage induced by platinum-based chemotherapy reagents, such as carboplatin^23^. As we observed that DPYSL3B depletion leads to downregulation of FA pathway members, we wanted to see if same effect is seen when DPYSL3B silencing is combined with FA pathway activating carboplatin. For this, we analysed the mRNA expression of key FA pathway members in PC-3 and DU-145 cells exposed to siDPYSL3B with or without carboplatin (DMSO ctrl) using RT-qPCR. Cells exposed to non-targeting siRNA (siCTRL) were used as control. Initially, we used 5 µM carboplatin concentration for both PC-3 and DU-145 cell lines and saw an upregulation of FA pathway members in response to carboplatin in siCTRL PC-3 cells (Fig. 3d). As expected, this upregulation was diminished upon depletion of DPYSL3B (Fig. 3d). DU-145 cells, however, appeared to lack a response (Supplementary Fig. S3a), which is why we decided to use 10 µM final carboplatin concentration instead. Increasing carboplatin concentration resulted in similar upregulation of FA pathway members with siCTRL in DU-145 cells (Fig. 3e) as was seen in PC-3 cells. Additionally, FA members FANCI and FANCD2, among others, were downregulated with the combined carboplatin and DPYSL3B silencing exposure when compared to cells exposed to carboplatin together with control siRNA (Fig. 3e).

Lastly, we wanted to investigate the ubiquitination status of FANCI using western blot, as the activation of FA pathway requires mono-ubiquitinated form of FANCI (mUB-FANCI). To this end, we exposed DPYSL3 silenced PC-3 and DU-145 cells to either DMSO or carboplatin using non-targeting siRNA exposed cells as control. Indeed, an induction of mUB-FANCI was observed in siCTRL cells in response to carboplatin in both PC-3 and DU-145 cell lines (Fig. 3f and g, respectively). However, we also noticed a difference between PC-3 and DU-145 in response to DPYSL3B depletion: while in PC-3 cells DPYSL3B silencing led to overall reduction in both FANCI and mUB-FANCI protein levels, in DU-145 cells, DPYSL3B silencing alone (siDPYSL3B+DMSO) promoted ubiquitination and thus activation of FANCI like exposure to carboplatin alone (siCTRL+carboplatin) (Fig. 3g). The combined carboplatin exposure with DPYSL3B depletion, in contrast, led to significant reduction in FANCI expression overall in DU-145 cells as neither FANCI nor its ubiquitinated form was detected (Fig. 3f and g). Taken together, these results reveal that DPYSL3B silencing affects FANCI and thereby the activation of the FA DNA repair pathway in PC-3 and DU-145 prostate cancer cells.

### DPYSL3B depletion induces DNA damage accumulation in prostate cancer cells

Since we detected downregulation of FA pathway members in response to DPYSL3B depletion, which in turn indicates diminished capacity to repair especially carboplatin induced DNA damage, we studied whether DPYSL3B depletion leads to accumulation of DNA damage. Thus, we performed immunofluorescence analysis with γH2AX as a marker of DNA damage to analyze the effects of DPYSL3B depletion with and without carboplatin in PC-3 and DU-145 cells. In both cell lines, DPYSL3B silencing alone increased the amount of γH2AX foci per nuclei when compared to control siRNA cells, thus indicating increased DNA damage (Fig. 4a and b). The effect was more pronounced in PC-3 cells (Fig. 4a) which coincides with our GSEA results showing negative enrichment of DNA damage repair related pathways in PC-3 cells but not in DU-145 cells.

**Figure 4.**
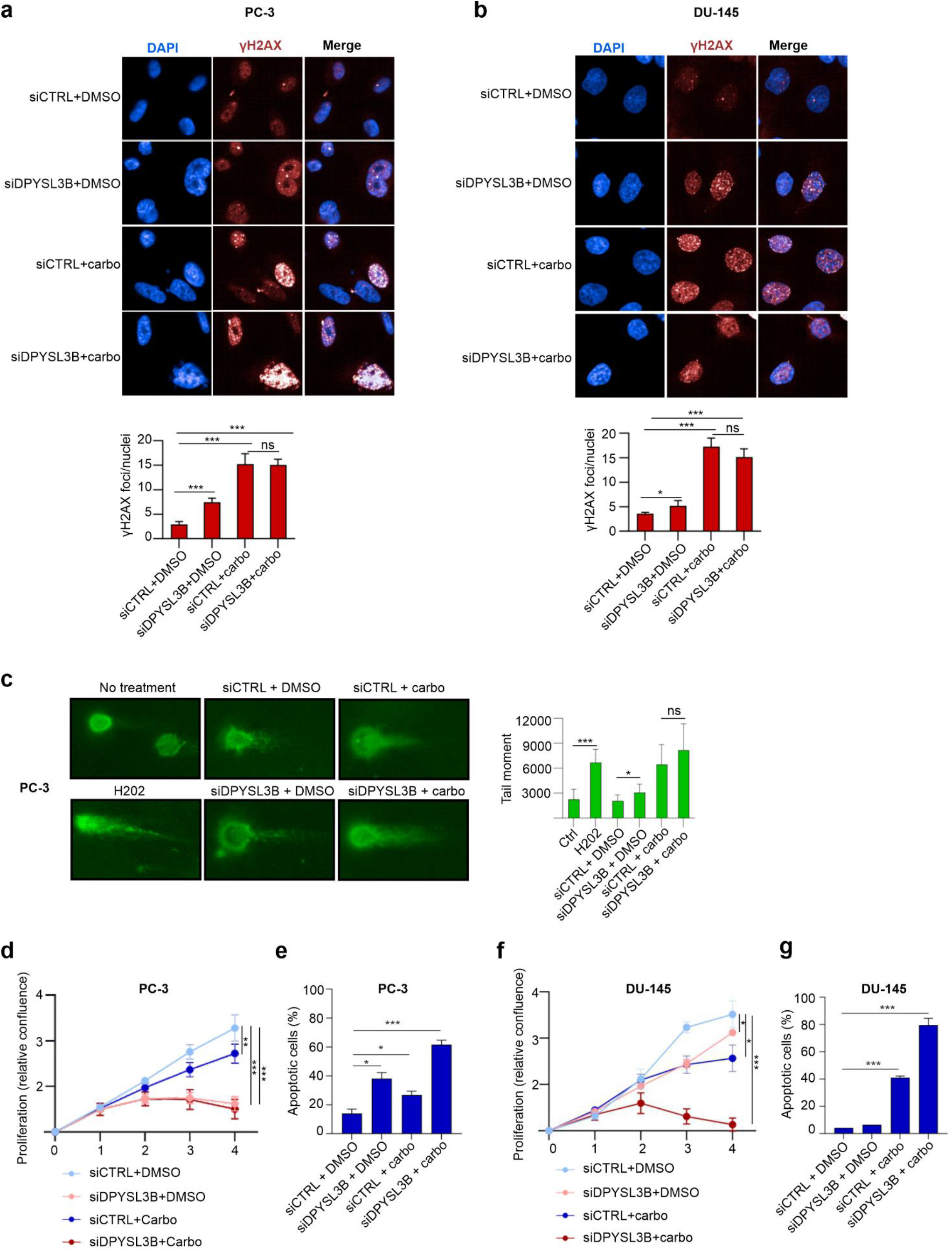
DPYSL3B depletion induces DNA damage accumulation and enhances sensitivity to carboplatin in prostate cancer cells. DNA damage was evaluated using immunofluorescence imaging and phosphorylated Histone 2AX (γH2AX) as DNA damage marker. Immunofluorescence images of a) PC-3 cells and b) DU-145 cells exposed to either siCTRL or siDPYSL3B in combination with DMSO or DNA damage inducing carboplatin stained with γH2AX (red) and DAPI (blue). In both cell lines DPYSL3B silencing alone (siDPYSL3B+DMSO sample) was shown to cause accumulation of DNA damage based on increased γH2AX signal when compared to control (siCTRL+DMSO). Addition of carboplatin resulted in significant increase in DNA damage when compared to DMSO exposed cells. Bars represent mean ± SD with n = 4 (individual treatments). p-values shown as asterisks (*p ≤ 0.05, **p ≤ 0.01 and ***p ≤ 0.001 c) DNA damage was also analysed using single cell electrophoresis assay known as comet assay. PC-3 cells exposed to siCTRL or siDPYSL3B together with DMSO or carboplatin were included in the analysis. Cells exposed to highly damaging hydrogen peroxide (H202) were used as positive control while untreated cells were used as negative control. The level of DNA damage is estimated based on the tail moment, which is calculated by multiplying the length of the DNA tail protruding from the nucleus with the percentage of DNA in the tail. DPYSL3B silencing alone was found to increase DNA damage. Also, in cells exposed to the combination carboplatin and DPYSL3B depletion, an increase in the tail moment was observed indicating DNA damage. Bars represent mean ± SD with n = 8. p-values shown as asterisks (*p ≤ 0.05, **p ≤ 0.01 and ***p ≤ 0.001 d) The effect of combined exposure to DPYSL3B siRNA and carboplatin on PC-3 cell proliferation was studied using IncuCyte live cell imaging system and the associated software. A significant decrease in proliferation was detected in response to siDPYSL3B alone (siDPYSL3B+DMSO, light red curve) and combination of both siDPYSL3B and carboplatin (siDPYSL3B+carboplatin, dark red curve) when compared to the corresponding controls (siCTRL+DMSO, light blue curve and siCTRL+carboplatin, dark blue curve). Curves represent mean ± SD with n = 4. p-values shown as asterisks (*p ≤ 0.05, **p ≤ 0.01 and ***p ≤ 0.001 e) The percentage of apoptotic cells in response to DPYSL3B siRNA alone (siDPYSL3B+DMSO) and in combination with carboplatin (siDPYSL3B+carboplatin) in PC-3 cells were quantified using Incucyte® Caspase-3/7 Red Dye and IncuCyte live-cell imaging system. PC-3 siCTRL cells with DMSO (siCTRL+DMSO) or with carboplatin (siCTRL+carboplatin) were used as comparison. Based on the results, DPYSL3B depletion alone induced apoptosis. The effect was more pronounced when siDPYSL3B was combined with carboplatin. Bars represent mean ± SD with n = 3. p-values shown as asterisks (*p ≤ 0.05, **p ≤ 0.01 and ***p ≤ 0.001 f) The proliferation of DU-145 cells exposed to either siCTRL+DMSO (light blue), siDPYSL3B+DMSO (light red), siCTRL+carboplatin (dark blue) or siDPYSL3B+carboplatin (dark red) was analysed using IncuCyte live-cell imaging in a similar manner as for PC-3 cells. A significant decrease in proliferation was detected in response to both siDPYSL3B alone (siDPYSL3B+DMSO) and to the combination of siDPYSL3B and carboplatin. g) The effect of DPYSL3B depletion alone (siDPYSL3B+DMSO) and in combination with carboplatin (siDPYSL3B+carboplatin) on apoptosis in DU-145 cells was analysed in a similar manner as for PC-3 cells. DPYSL3B silencing caused only minor increase in apoptotic cells when compared to control siRNA (siCTRL+DMSO) while majority of the cells were apoptotic when exposed to the combination of siDPYSL3 with carboplatin. Results are shown as a bar graph with percentages of apoptotic DU-145 cells.

Exposure to carboplatin dramatically increased DNA damage in both PC-3 and DU-145 cell lines when compared to DMSO exposed cells based on the calculated number of γH2AX foci per nuclei (Fig. 4a and b). However, no additive effect was observed when combining carboplatin with DPYSL3B depletion. To further confirm our observations, we performed comet assay to assess DNA damage in PC-3 cells in response to DPYSL3B silencing alone and in combination with carboplatin. The results showed an increase in tail moment (length of the DNA tail relative to untreated cells), an indicator of DNA damage, in DPYSL3B depleted cells when compared to siCTRL (Fig. 4c). When comparing carboplatin exposed siDPYSL3B and siCTRL cells, no additive effect on DNA damage was observed based on the calculated tail moment. Overall, our results show that DPYSL3 depletion alone results in accumulation of DNA damage loci in prostate cancer cells but does not potentiate the carboplatin-induced DNA damage.

### DPYSL3B depletion promotes apoptosis and sensitizes prostate cancer cells to carboplatin

Carboplatin is commonly used in the treatment of AR-negative PCa but resistance often emerges. Earlier, we showed that DPYSL3B silencing alone reduces proliferation in PC-3 and DU-145 cells (Fig. 2a-c). We therefore hypothesized that DPYSL3B depletion may sensitize PCa cells, particularly AR-negative PC-3 and DU-145 cells, to carboplatin-induced apoptosis. First, we assessed the effect of carboplatin alone (siCTRL+DMSO vs siCTRL+carboplatin) in PC-3 cells and observed a reduction in proliferation (Fig. 4d). However, when DPYSL3-silenced cells (siDPYSL3B) were compared to those exposed to both DPYSL3 siRNA and carboplatin, no significant additive effect on proliferation was detected, likely because DPYSL3 depletion alone caused a profound growth arrest (Fig. 4d). Live-cell imaging suggested increased apoptosis in siDPYSL3 cells, with or without carboplatin, prompting further validation (Supplementary Fig. S4a). This prompted us to further investigate the effect of DPYSL3 depletion on apoptosis in PC-3 cells using a live-cell caspase-3/7 activity assay. In PC-3 cells approximately 40 % of the cells were apoptotic in response to DPYSL3B depletion alone, compared to siCTRL (Fig. 4d, bar graph). When combined with carboplatin, apoptosis increased to 60%, significantly higher than with either DPYSL3B depletion or carboplatin alone (the latter ∼30%) (Fig. 4d, bar graph).

Interestingly, in DU-145 cells, DPYSL3B silencing alone did not show as dramatic effect on proliferation or on apoptosis as seen in PC-3 cells (Fig. 4e). However, when carboplatin was included, proliferation was significantly reduced both with control and DPYSL3 siRNA (siCTRL+carboplatin and siDPYSL3+carboplatin) in comparison to the corresponding DMSO controls (Fig. 4e). The effect of carboplatin was significantly more pronounced in DPYSL3B silenced cells as the overwhelming majority (80 %) of the cells were apoptotic (Fig. 4e, bar graph).

Overall, the results indicate that DPYSL3B is required for prostate cancer cell survival likely through its regulatory role in DNA damage repair.

## Discussion

Treatment of CRPC subtypes that do not respond to AR inhibition, such as DNPC and t-NEPC, relies primarily on taxanes or platinum-based chemotherapies^42^. Patients harboring germline mutations in DNA damage response genes, such as *BRCA1* (*FANCS*) and *BRCA2* (*FANCD1*), have been shown to benefit most from these therapies^43^. However, resistance to chemotherapy frequently occurs through mechanisms such as increased drug efflux, enhanced DNA repair, or glutathione-mediated detoxification^44^. Therefore, chemotherapy resistance remains a major clinical challenge.

In the present study, we characterized the role of DPYSL3B in prostate cancer and found it to be highly upregulated in DNPC tumors as well as in AR-negative cell lines. Functional depletion of DPYSL3B resulted in disrupted cellular metabolism, reduced proliferation, and downregulation of Fanconi anemia complex, particularly FANCI. This was accompanied by accumulation of DNA damage, as evidenced by increased γH2AX levels and comet assay analysis. Furthermore, DPYSL3B depletion sensitized prostate cancer cells to the DNA-damaging agent carboplatin and led to apoptosis in DU-145 and PC-3 cells. Consistently, high DPYSL3B expression was associated with poor overall survival in prostate cancer patients based on transcriptomic data, suggesting a potential clinical relevance for DPYSL3B in aggressive disease.

Previous research has shown that the DPYSL3A isoform undergoes N-terminal cleavage facilitated by calpain in PC-3 cells^20^. Calpain is a calcium-dependent protease which can be activated by increased Ca^2+^ influx induced by ROS^45^. The N-terminal part of DPYSL3A is able to translocate to the nucleus to the promoter region of E2F1, enhancing its expression. Thus, we propose that DPYSL3B could undergo similar cleavage, as both DPYSL3A and B share the cleavage site in their N-terminal part. To further support this hypothesis, we detected downregulation of E2F1 as well as its targets. Interestingly, E2F1/2 has been shown to regulate expression of members of FA-complex^46,47^. We therefore propose a DPYSL3B-E2F1-FA axis as a cellular mechanism to induce DNA repair in response to oxidative stress (Fig. 5).

**Figure 5.**
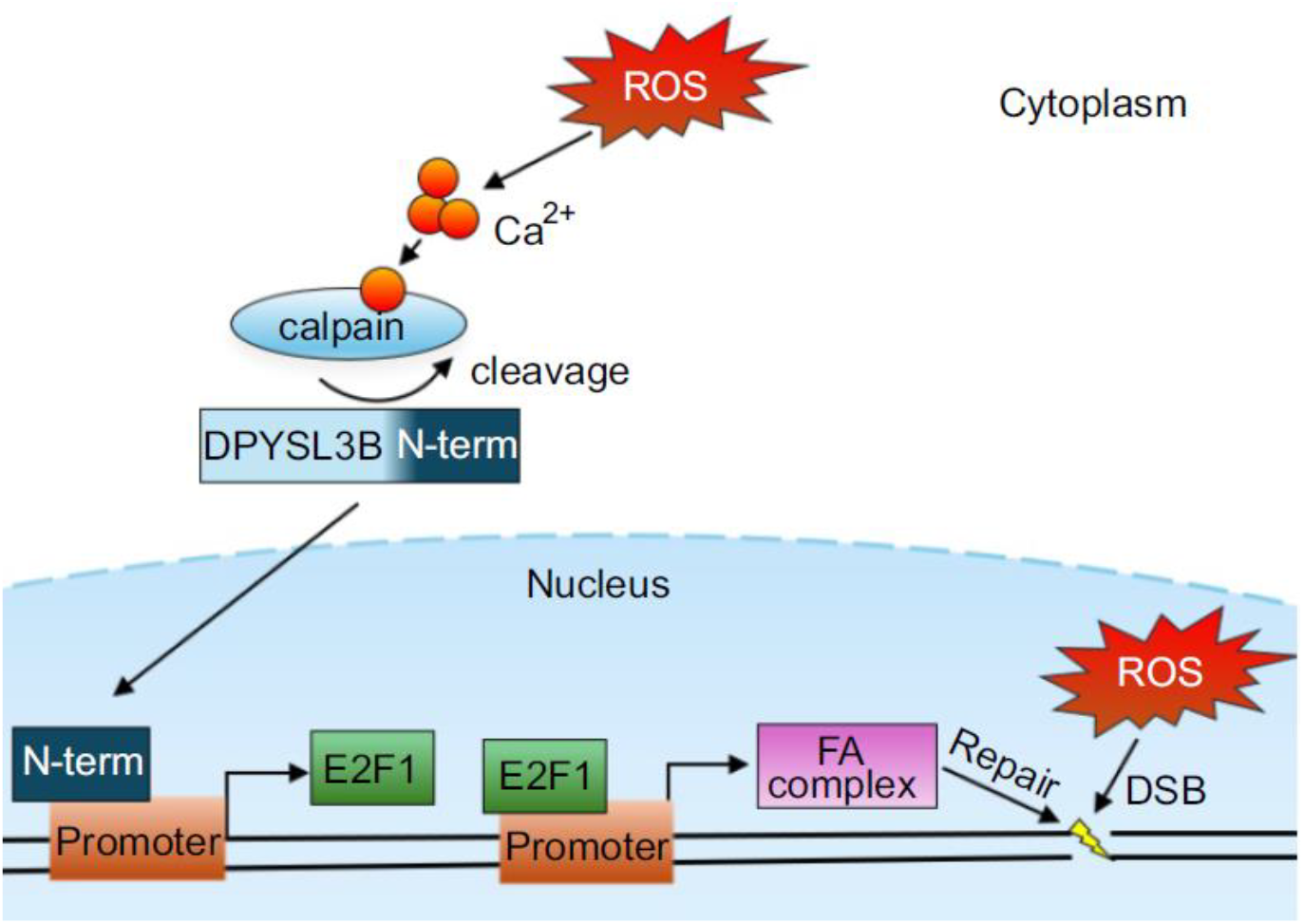
Proposed mechanism of DPYSL3B function. Reactive oxygen species (ROS) induce calcium influx, leading to elevated intracellular calcium levels. This increase in calcium activates calpain, a calcium-dependent, non-lysosomal cysteine protease. Activated calpain cleaves DPYSL3B and induces nuclear translocation of the cleaved N-terminal fragment. In the nucleus, the N-terminal part of DPYSL3B binds to the promoter region of *E2F1*, thereby enhancing its transcription. *E2F1*, in turn, increases the expression of FA pathway members, promoting DNA repair in response to ROS-induced damage.

Additionally, we detected differential responses to DPYSL3B depletion between PC-3 and DU-145 cells. PC-3 cells were more sensitive, showing stronger inhibition of proliferation and greater downregulation of FANCI at both the mRNA and protein levels. One possible explanation for this difference could be in the ability of DPYSL3B to mediate its effects via E2F1 as it has been shown that inhibition of both E2F1/E2F2 induces DNA damage in both PC-3 and DU-145 cells but reduces cell viability more in PC-3 than DU-145 cell^48^. This is in line with our observations on the effects of DPYSL3B depletion. Additionally, in another study that compared the effect of E2F1 depletion alone in PC-3 and DU-145 cells, it was discovered that E2F1 knockdown in PC-3 cells led to reduced proliferation and extensive changes in transcriptome, whereas in DU-145 cells the effect was non-existent^49^. To summarize, DU-145 cells may be less dependent on E2F1 and more reliant on E2F2 for FA pathway gene regulation. Since DPYSL3B is hypothesized to act via E2F1, this could partially explain why FANCI downregulation was not observed in response to DPYSL3B depletion alone.

Based on our RNA-seq and metabolomics analysis data, DPYSL3B may have other E2F-independent functions. Previously it has been shown that DPYSL3B depletion leads to downregulation of glucose transporter GLUT1 and hinders mTOR signaling in urothelial carcinoma^50^. Similarly, we also detected that DPYSL3B depletion led to reduced levels of both metabolites required for glutathione synthesis and glutathione itself. Glutathione is an important antioxidant protecting cells from oxidative damage and its metabolism involves glycine, cysteine and glutamic acid. We found that DPYSL3B depletion resulted in a reduction in glutamic acid levels accompanied by accumulation of L-cystine, which is an oxidized form of cysteine. This is an indication of an increase in oxidative stress in response to DPYSL3B depletion. Interestingly, it has been shown that glucose deprivation induced by GLUT1 inhibition increases l-cystine import^51^. Furthermore, transformation of l-cystine into cysteine for GSH production requires NAPDH which is quickly depleted in high l-cystine environment^51^. The depletion of both GSH and NAPDH leads to accumulation of ROS. Our hypothesis is that DPYSL3B depletion increases l-cystine intake leading to consumption of NAPDH and eventual failure to regulate ROS, which in turn leads to induction of DNA damage. This is supported by our metabolomics analysis and DNA damage assays on DPYSL3B depleted PC-3 and DU-145 cells. Although further mechanistic validation is needed, our metabolomic and transcriptomic data support a model in which DPYSL3B depletion disrupts antioxidant homeostasis and contributes to DNA damage accumulation through elevated oxidative stress.

We discovered that DU-145 cells are less sensitive to carboplatin than PC-3 cells. In PC-3 cells lower carboplatin concentration was sufficient to induce DNA damage and expression of FA-pathway members when compared to DU-145 cells. This observation is in line with previous research, as DU-145 cells have been shown to be more resistant than PC-3 cells towards DNA damage inducing treatment, such as IR and cisplatin^52,53^. One potential explanation for the higher carboplatin tolerance could be Keap1 inactivation as a result of Keap1 promoter hypermethylation found in DU-145 cells but not in PC-3 cells^54^. Keap1 is an inhibitor protein of Nrf2, which in turn regulates the expression of antioxidant proteins and protects the cells against oxidative damage caused by radiation-and platinum-based therapies^54^. Therefore, DU-145 cells could be more tolerant towards oxidative stress induced by DPYSL3B depletion than PC-3 cells due to the inactivation of Keap1 and subsequent expression of Nrf2. It should be noted that we did observe mUB-FANCI in response to DPYSL3B depletion alone in DU-145 indicating activation of FA-pathway in response to DPYSL3B depletion. The expression of FANCI was diminished upon combined exposure to DPYSL3B depletion and carboplatin, leading to induction of apoptosis. Therefore, DPYSL3B silencing was shown to sensitize also DU-145 cells to carboplatin.

In conclusion, our results suggest that DPYSL3B is involved in DNA damage response via the FA pathway, and that its depletion sensitizes prostate cancer cells to DNA-damaging chemotherapy. These findings identify DPYSL3B as a potential therapeutic target to overcome resistance to platinum-based therapies and highlight its possible role in DNA repair and metabolic reprogramming, notably affecting redox balance and biosynthetic regulation, particularly in AR-negative and double-negative prostate cancer, and provide a foundation for further studies to explore its potential in clinical applications.

## Materials and methods

### In silico analyses

Median RNA-seq RNA expression values and clinicopathological data from different prostate cancer datasets were analysed using cBioPortal^55,56^.

### Cell culture

LNCaP, PC-3 and DU-145 cells were cultured in RPMI 1640 media (Gibco) supplemented with 10% FBS (Thermo Fisher Scientific), 2 mM L-glutamine (Thermo Fisher Scientific) and streptomycin/penicillin.

### siRNA transfections

Cells were reverse transfected with 25 nM of DPYSL3B sirna or 25 nM of non-targeting siRNA (siCTRL) (ON-TARGETplus SMARTpool siRNA, Dharmacon) using OPTI-MEM and RNAiMAX (Invitrogen). Cells were cultured without streptomycin and penicillin to maximize transfection efficiency.

### Proliferation and apoptosis analysis

Cells (5 000 cells per well for LNCaPs, 3000 for PC-3, 2 500 for DU-145) were transfected with siRNA and seeded on 96-well plates. After 24 h, DMSO or carboplatin (Selleck Chemicals GmbH) were added on plates. Plates were imaged daily, and confluence was analysed with IncuCyte S3 Live-Cell software. Apoptotic cells were determined by using Caspase-3/7 reagent (Sartorius, red for DU-145 cells and green for PC-3 cells) and fluorescent signal was measured with IncuCyte S3 Live-Cell software.

### Quantitative PCR and RNA-sequencing

Cells (300 000 LNCaPs, 150 000 PC-3 and DU-145) were plated on 6-well plates and transfected with siRNA. After 24 h, carboplatin or DMSO was added. After 72 h from transfection, samples (n=3) were collected. RNA was extracted from cells with TriPure Isolation Reagent (Invitrogen). For qPCR, RNA was reverse transcribed with Transcriptor First Strand cDNA synthesis Kit (Roche). Gene expression was analysed using LightCycler 480 SYBR Green I Master (Roche), LightCycler 480 II (Roche) and 2-ΔΔCt method. For RNA-sequencing, DPYSL3B siRNA and non-tageting siRNA were sent to Novogene (Cambridge, UK) for quality check, library preparation and sequencing. Analysis was performed with Nextflow pipeline^57^.

### Targeted metabolomics profiling analysis

Cells (300 000 LNCaPs, 150 000 PC-3) were plated on 6-well plates and transfected with siRNA. After 72 hours from transfection, cells were washed with ice cold PBS, scared into ice cold PBS, pelleted with centrifuge and stored in -20°C.

The metabolites were extracted using cold extraction solvent (Acetonitrile:Methanol:Milli-Q Water; 40:40:20, LS-MS grade, Thermo Fischer Scientific) followed by sonication for 60 seconds (power 60 and frequency 37), vortexing for three cycles and centrifugation at 14000rpm at 4°C for 5 minutes. The supernatants were evaporated to dry under Nitrogen gas. Dried samples were resuspended into extraction solvent with vortexing. Finally, the samples were injected randomized into a Thermo Vanquish UHPLC system coupled to a Q-Exactive Orbitrap quadrupole mass spectrometer (MS) equipped with a heated electrospray ionization (H-ESI) source probe (Thermo Fischer Scientific). Chromatographic separation was performed with a SeQuant ZIC-pHILIC column (2.1×100 mm, 5-μm particle; Merck), flow rate of 0.100 ml/minutes and mobile phase gradient with 20mM ammonium hydrogen carbonate, adjusted to pH 9.4 with ammonium solution (25%) as mobile phase A and Acetonitrile as mobile phase B, 0-2 min 80 % B, 2-17 min 80-20 % B, 17-24 min 80 % B. The column oven and auto-sampler temperatures were set to 40 ± 3 °C and 5 ± 3 °C, respectively. Scanning was performed with MS1 mode, mass range from 55 to 825 m/z using polarity switching and the following settings: resolution of 70,000, spray voltages of 4250 V for positive mode and 3250 V for negative mode, sheath gas flow rate of 25 arbitrary units (AU), auxiliary gas flow rate of 15 AU, sweep gas flow rate of 0, capillary temperature of 275°C, and S-lens RF level of 50.0. The instrument was controlled with the Xcalibur 4.1.31.9 software (Thermo Fischer Scientific). The metabolite annotations and integrations were done with the TraceFinder 5.1 software (Thermo Fischer Scientific) using confirmed retention times by in-house standard library (MSMLS-1EA, Merck) and compound m/z. The instrument operation was monitored throughout the run using pooled Quality Control (QC) sample prepared by pooling 5µL from each suspended sample and interspersed throughout the run as every 10th sample with the blank (extraction solvent). The data processing was monitored for peak quality (poor chromatography) and RSD > 20 % of pooled QC and background % >20 % noise (mean pooled QC/blank) and noted down.

### Gene set enrichment analysis

Gene expression datasets were subjected to gene set enrichment analysis (GSEA) using GSEA software (v.4.3.3) from the Broad Institute (Massachusetts Institute of Technology)^58^. The GSEA run was performed in weighted mode to identify significantly enriched pathways. Pathways enriched with a nominal P-value <.05 and false discovery rate (FDR) <0.25 were considered as significant.

### Western Blot

Cells (300 000 LNCaPs, 150 000 PC-3 and DU-145) were plated on 6-well plates and transfected with siRNA. After 24 h, carboplatin or DMSO was added. After 72 h from transfection, samples (n=2) were collected. Cell lysates were prepared with SDS buffer containing protease inhibitor (cOmplete™ Protease Inhibitor Cocktail, Roche) and cells were lysed with sonication. Proteins were separated with SDS-PAGE and transferred to nitrocellulose membranes. Membranes were blocked for 1 hour in 5 % non-fat dry milk (Valio) in 1xTBS-Tween. Primary antibodies were incubated in 4°C overnight and HRP-conjugated secondary antibodies were incubated for 1 h in RT. Proteins were detected Pierce™ ECL Western Blotting Substrate (Thermo Fisher Scientific), and membranes were imaged with Chemidoc (Bio-Rad).

### Immunofluorescence imaging

Cells (3 000 PC-3 or DU-145 cells) were seeded on PhenoPlate 96-well plate and transfected with siRNA. After 24 h carboplatin or DMSO was added. 3 days from transfection, cells were fixed with 4% paraformaldehyde, permeabilized with 0,1 % Triton-X, blocked 30 minutes with 1 % BSA in RT and incubated overnight in 4°C with primary antibody of γH2AX (Cell Signaling Technology) with 1:150 dilution. The next day, secondary antibody (Alexa Fluor™ 633 Goat anti-Rabbit IgG (H+L)), was added with DAPI (Sigma) and incubated in RT for 1 hour. Plates were imaged with Opera Phenix (PerkinElmer), and Harmony High-content imaging and analysis software was used to analyze the images.

### Neutral comet assay

PC-3 cells were seeded on 6-well plates and transfected with siRNA and after 24h from transfection, carboplatin or DMSO was added. After 72 h from transfection, hydrogen peroxide was added in 1:100 to create positive control samples. A neutral comet assay was performed according to manufacturer’s information (Bio-techne). DNA was stained with SYBR Gold (Cat.no S11494, Invitrogen) and imaged with confocal microscope (Zeiss LSM 700, Carl Zeiss Microimaging GmbH).

## Acknowledgements

We thank Onni Niemi and Merja Räsänen for excellent technical assistance. This work was carried out with the support of UEF Cell and Tissue Imaging Unit, University of Eastern Finland, Biocenter Kuopio and Euro-BioImaging Finland. The computational analyses were performed on servers provided by UEF Bioinformatics Center, University of Eastern Finland, Biocenter Finland, Finland. The facilities and expertise of Helsinki Metabolomics Center, supported by the Faculty of Medicine/University of Helsinki and HiLIFE is gratefully acknowledged.

## Author contributions

RK: Conceptualization, Methodology, Validation, Formal analysis, Investigation, Writing - Original Draft, Visualization, Project administration, Data Curation. HK: Conceptualization, Methodology, Validation, Formal analysis, Investigation, Writing - Original Draft, Data Curation, Project administration. NI: Formal analysis, Investigation, Data Curation. KB: Formal analysis, Software, Investigation, Data Curation. KK: Conceptualization, Investigation, Funding acquisition, Project administration, Supervision, Writing - Original Draft, Resources.

