## Supplementary file S1 for "DPYSL3B is a regulator of chemoresistance via DNA repair and metabolic reprogramming in prostate cancer"

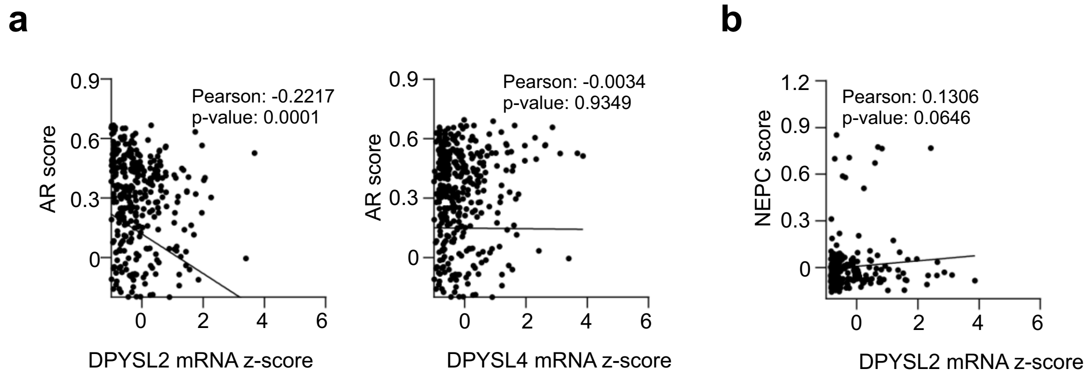


**Supplementary Figure 1.** a) Correlation between AR score and DPYSL2 and DPYSL4 mRNA z-score. DPYSL2 expression negatively correlates with AR score b) Correlation between NEPC score and DPYSL2 and DPYSL4 mRNA z-score. No correlation is detected
