## Supplementary file S2 for "DPYSL3B is a regulator of chemoresistance via DNA repair and metabolic reprogramming in prostate cancer"

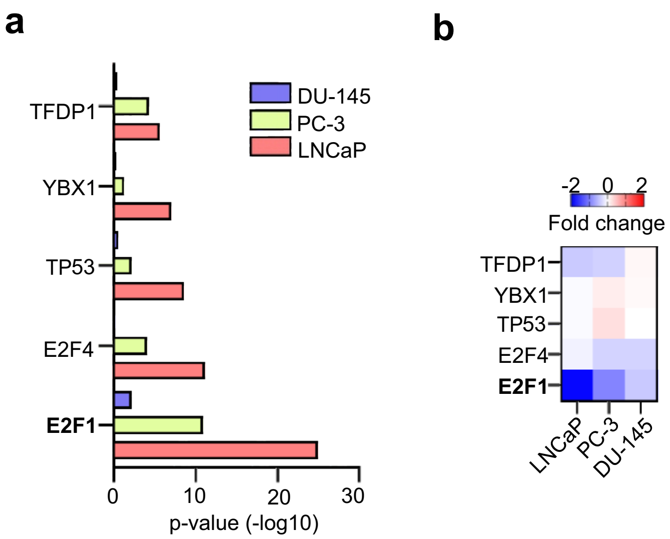


**Supplementary figure 2.** a) Transcription factor enrichment analysis of downregulated differentially expressed genes in DU-145, PC-3 and LNCaP cells. Downregulated DEGs in all cell lines show enrichment with E2F1 targets. b) Expression of transcription factors in LNCaP, PC-3 and DU-145 cells. Expression of E2F1 is downregulated in all cell lines.
