## Supplementary file S3 for "DPYSL3B is a regulator of chemoresistance via DNA repair and metabolic reprogramming in prostate cancer"

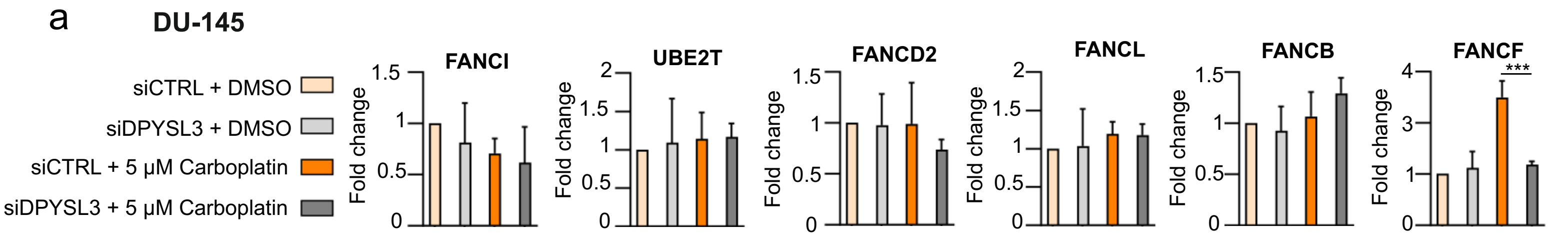


**Supplementary figure 3** a) Carboplatin or DPYSL3B depletion doesn't affect mRNA expression of FANCI, UBE2T, FANCD2, FANCL, FANCB. FANCF expression is induced in respons to Carboplatin and is downregulated when combined with DPYSL3B siRNA
