## Supplementary file S4 for "DPYSL3B is a regulator of chemoresistance via DNA repair and metabolic reprogramming in prostate cancer"

a

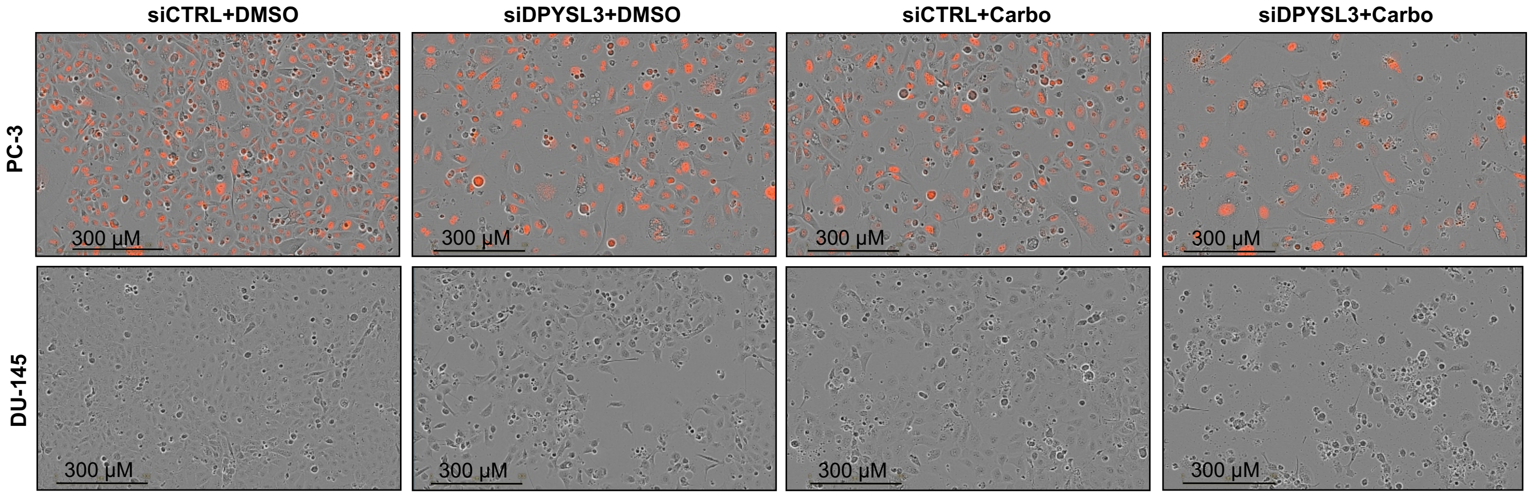


**Supplementary Figure 4.** a) Images of PC-3 nucred cells and DU-145 cells. Cells appear apoptotic in response to combination of Carboplatin and DPYSL3B depletion (last panel).
